# MiR-146a controls age related bone loss

**DOI:** 10.1101/2020.03.20.000174

**Authors:** Victoria Saferding, Melanie Hofmann, Julia S. Brunner, Birgit Niederreiter, Melanie Timmen, Nathaniel Magilnick, Silvia Hayer, Gerwin Heller, Günter Steiner, Richard Stange, Mark Boldin, Gernot Schabbauer, Moritz Weigl, Matthias Hackl, Johannes Grillari, Josef S. Smolen, Stephan Blüml

## Abstract

Bone loss is one of the consequences of aging, leading to diseases such as osteoporosis and increased susceptibility to fragility fractures and therefore considerable morbidity and mortality in humans. Here we identify microRNA 146a as an essential epigenetic switch controlling bone loss with age. Mice deficient in miR-146a show regular development of their skeleton. However, while WT mice start to lose bone with age, animals deficient in miR-146a continue to accrue bone throughout their life span. Increased bone mass is due to increased generation and activity of osteoblasts in miR-146a deficient mice as a result of sustained activation of bone anabolic Wnt signaling during aging. Deregulation of the miR-146a target genes Wnt1 and Wnt5a parallel bone accrual and osteoblast generation, which is accompanied by reduced development of bone marrow adiposity. Furthermore, miR-146a deficient mice are protected from ovariectomy induced bone loss. In humans, levels of miR-146a are increased in patients suffering fragility fractures in comparison to those who do not.

These data identify miR-146a as a crucial epigenetic temporal regulator which essentially controls bone homeostasis during aging by regulating bone anabolic Wnt signaling. Therefore, miR-146a might be a powerful therapeutic target to prevent age related bone dysfunction such as the development of bone marrow adiposity and osteoporosis.

## Introduction

As bone is continuously remodeled, bone mass of vertebrates is constantly changing and determined by the balance of bone resorption and bone formation. In adolescence, bone formation predominates until the peak bone mass is reached. From then on, bone mass continuously declines, with certain events such as the hormonal changes in female menopause, accelerating the process [1]. Reduced bone mass is associated with reduced stability of bones in humans, leading to pathological conditions such as fragility fractures, which are a significant health problem, especially as life expectancy is constantly increasing [2, 3]. Bone loss can be caused by reduced bone formation as well as increased bone resorption, and many factors contributing to bone loss have been identified [2, 4].

MiR-146a is a predominantly anti-inflammatory microRNA that has been shown to be important in various aspects of inflammation and immunity [5, 6]. It has primarily been reported as an anti-inflammatory miRNA, as deficiency in miR-146a results in increased inflammation in various instances [7–9]. This miRNA has also been shown to be associated with aging, as it is one of the microRNAs that was demonstrated to increase with age [10], while it is reduced by hormone replacement therapy in post-menopausal women [11].

Several miRNAs, among them miR-146a, have been implicated in regulating the biology of bone, both osteoblasts/osteocytes and osteoclasts [12–17]. We have previously demonstrated that loss of miR-146a increases the severity of inflammatory arthritis by regulating fibroblast pathogenicity, especially their ability to induced bone resorbing osteoclasts [8]. However, the role of miR-146a in bone homeostasis is not known. Here we describe the microRNA-146a as a critical factor, which regulates bone loss during aging. Mice deficient in this miRNA constantly accrue bone over time, leading to a high bone mass phenotype. Loss of miR-146a also protects from ovariectomy induced bone loss. In addition, in humans we find increased levels of miR-146a in patients suffering fragility fractures compared to those who had no fractures, suggesting a crucial role of miR146a in bone homeostasis.

## Results

### MiR-146a deficient mice continuously accumulate bone during aging

To investigate the impact of miR-146a on bone biology we used miR-146a full knock out animals and assessed trabecular and cortical bone parameters in tibial bones of animals aged 3 to 16 months. Micro computed tomographic (μCT) analysis showed regular development of the skeleton and bone volume until 4 months of age until reaching the peak bone mass, which is at 3-4 months in C57BL/6 mice [18]. While WT mice started to lose bone mass from then on, as expected, miR-146a deficient mice continued to accrue bone until 12 months of age, with bone mass remaining stable until 16 months of age (Fig. 1 A-G). MiR-146a deficient mice displayed elevated trabecular bone volume per tissue volume (BV/TV), trabecular number (Tb.N), connectivity density (Conn. D) and reduced trabecular separation (Tb.Sp) compared to WT mice starting at 6 months of age throughout 16 months, while trabecular thickness (Tb. Th) was not changed between these two groups. Moreover, the structural model index (SMI) was significantly changed in miR-146a deficient animals, indicating a rather plate like than rod-like geometry of trabecular bone in the absence of miR-146a. In addition to trabecular measures of bone volume, cortical BV/TV and cortical thickness (Ct.Th) also increased and cortical porosity decreased significantly with age in miR-146a^−/−^ compared to control animals (Fig. 1 I and J). Similar to the observations in trabecular bone, cortical bone parameters in both groups such as cortical thickness (Ct.Th), cortical BV/TV and cortical porosity were not different until 4 months of age. However, starting from 6 months of age, cortical thickness and BV/TV increased, and cortical porosity decreased significantly in miR-146a deficient mice compared to WT mice (Fig. 1 I-K). Bone mineral density (BMD) solely increased at 16 months of age in knock out animals (Fig. 1 L). Of note, aged miR-146a deficient animals show massive trabecular bone growth reaching into the diaphyseal shaft of the tibia, which is normally devoid of trabecular bone in WT animals, indicating a strongly deregulated bone growth in miR-146a^−/−^ animals (Fig. 1 M). Interestingly, this bone phenotype was predominantly found in female mice, whereas male mice showed only a slight increase in their bone mass at 12 months of age (Sup. Fig. 1).

**Fig.1.**
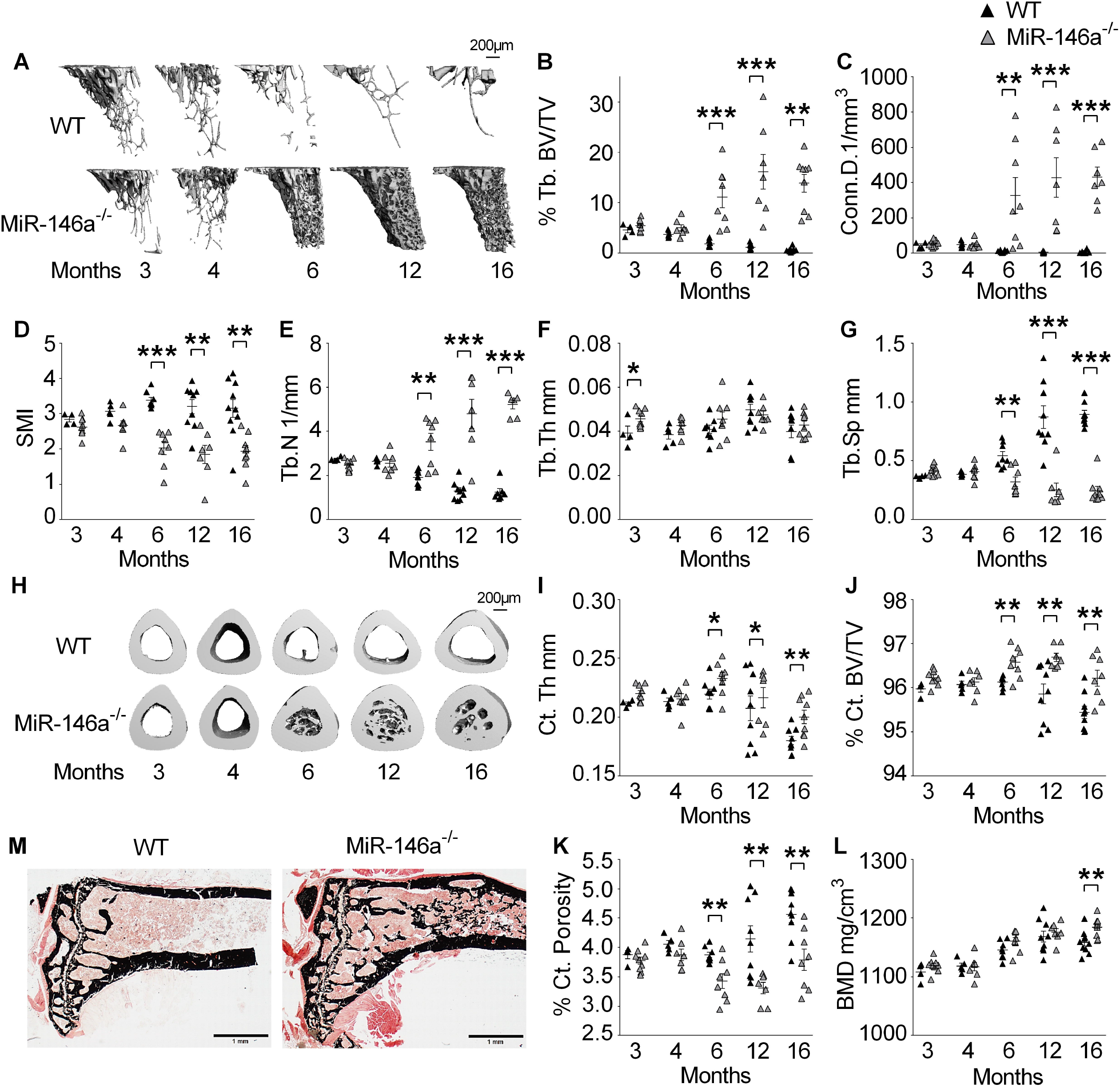
MiR-146a deficient mice continuously accumulate bone during aging. **A** Representative μCT images of trabecular bones from tibial WT and MiR-146a^−/−^ aged 3 to 16 months (bar 200μm). **B-G** Three-dimensional reconstruction and quantitation of the indicated parameters of trabecular bone from proximal tibias of WT and MiR-146a deficient animals aged 3 to 16 months, using μCT (n≥4). **H** Representative images of cortical bone from WT and MiR-146a^−/−^ animals aged 3 to 16 months (bar 200μm). **I-L** Bone morphometric analysis of the indicated parameters of cortical bone at the diaphysis of the tibia, close to site of the intersecting fibula using μCT(n≥4). **M** Von Kossa staining of histological sections, obtained from the proximal region of tibias from 6-month-old WT and MiR-146a^−/−^ mice, representative images are shown (bars 1mm, magnification x 2.5). Tb. BV/TV, trabecular bone volume per tissue volume; Conn. D, connectivity density; SMI, structural model index; Tb.N, trabecular number; Tb. Th, trabecular thickness; Tb. Sp, trabecular separation; Ct.Th, cortical thickness; Ct.BV/TV, cortical bone volume per tissue volume; BMD, bone mineral density; Ct. Porosity; cortical porosity. All data shown were obtained from female animals *p < 0.05; **p < 0.01; ***p < 0,001 Results are shown as mean ± SEM.

As the increase in bone mass was developing with age, we investigated, whether miR-146a levels were differentially regulated over time in bone of WT animals. Young and early adult mice at the age of 2 and 3 months showed no difference in the expression level of miR-146a in bone tissue (Fig. 2 A). However, starting at 4 months of age, expression levels of this micro RNA increased significantly, reaching a maximum at 5 months of age. Even though it declined from 5 to 6 months of age, its expression was still increased at this point, finally dropping in tissue samples of 12 months old animals (Fig. 2 A). These data demonstrate elevated miR-146a expression in bone around peak bone mass, and the course of miR-146a expression correlates with the switch from bone accrual to bone loss, supporting the notion of a regulatory function of miR-146a in bone during aging.

**Fig. 2.**
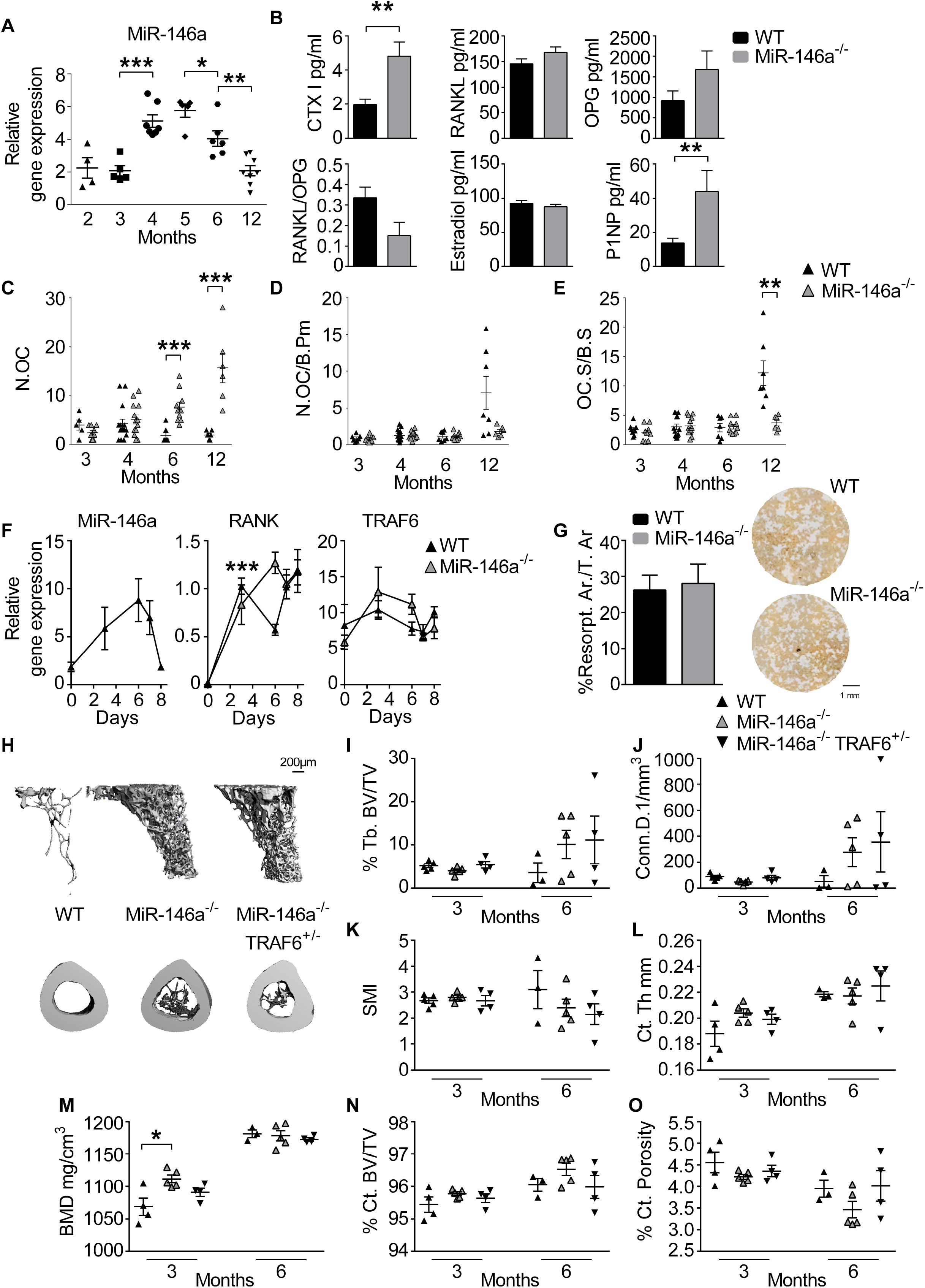
Osteoclast activity is not responsible for high bone mass in MiR-146a deficient animals. **A** Expression level of MiR-146a in femoral bone of WT animals aged from 2 to 12 months using quantitative real time PCR (Q-PCR) analysis (n≥4). **B** Levels of CTX, RANKL, OPG, Estradiol, P1NP and RANKL/OPG ratio were measured in sera of 6-month-old WT and MiR-146a^−/−^ mice using ELISA (n≥4). **C-E** Histomorphometric analysis of tartrate-resistant acid phosphatase (TRAP) stained tibial sections, N.OC, N.OC/B.Pm and OC.S/BS of WT and MiR-146a^−/−^ mice aged 3 to 12 months were assessed (n≥5). **F** Expression level of MiR-146a, RANK and TRAF6 in bone marrow derived osteoclasts from 3 months old WT and MiR-146a^−/−^ animals was analyzed using Q-PCR. Bone marrow was isolated on day 0, cells were cultured with MCSF over 8 days and stimulated with RANKL on day 3 and 6 (n=4). **G** Quantification of osteoclast resorption capacity (left) of WT and MiR-146a deficient animals. Representative images of *in vitro* bone resorption assays are shown (right, bar 1mm) (n=8). **H** Representative μCT images of trabecular and cortical bone of 6 months old WT, MiR-146a^−/−^ and MiR-146a^−/−^ TRAF6^+/−^ animals are shown (bar 200μm). **I-O** Bone morphometric analysis of trabecular bone from proximal tibias and cortical bone at the diaphysis of the tibia (close to site of the intersecting fibula) of 3- and 6-months old WT, MiR-146a^−/−^ and MiR-146a^−/−^ TRAF6^+/−^ mice (n≥3). N.OC, numbers of osteoclasts; N.OC/B.Pm, numbers of osteoclasts per bone perimeter; OC.S/BS, osteoclast surface per bone surface; CTX, C-terminal telopeptide of type I collagen; RANKL, Receptor Activator of NF-κB Ligand; OPG, osteoprotegerin; P1NP, procollagen type 1 N-terminal propeptide; Tb. BV/TV, trabecular bone volume per tissue volume; Conn. D, connectivity density; SMI, structural model index; Ct.Th, cortical thickness; BMD, bone mineral density; Ct.BV/TV, cortical bone volume per tissue volume; Ct. Porosity; cortical porosity; All analysis were performed in female animals *p < 0.05; **p < 0.01; ***p < 0,001 Results are shown as mean ± SEM.

Following, we examined known regulators of bone turnover in sera of 6 months old WT and miR-146a^−/−^ animals. MiR-146a^−/−^ animals showed elevated levels of the bone resorption marker C-terminal telopeptide of type I collagen (CTXI) as well as the bone formation marker P1NP. Since the bone phenotype was more profound in female animals, we assessed estrogen levels, which were not different between WT and knock out animals. Moreover, we did detect a trend towards a reduced RANKL/OPG ratio in miR-146a deficient mice compared to WT mice which was primarily driven by increased levels of OPG in miR-146a^−/−^ animals compared to WT animals (Fig. 2 B).

Analysis of marker genes of osteoblasts including Runt-related transcription factor 2 (Runx2), Osteocalcin (OC), Alkaline Phosphatase (ALP), Osterix (OSX), Osteopontin (OPN) as well as osteoclasts, including Tartrate resistant acid phosphatase (TRAP), Cathepsin K(CTSK) and Receptor Activator of NF-κB (RANK) in femoral bones of WT and miR-146a^−/−^ animals did not show any difference at 3 and 6 months of age. However, RUNX2, OSX, OPN as well as TRAP, CTSK and RANK increased significantly at 12 months of age in miR-146a deficient bones (Sup. Fig. 2 A and B). Taken together these data demonstrate, that both osteoblast as well as osteoclast associated genes are deregulated in miR-146a^−/−^ mice during aging.

Therefore, we investigated static histomorphometry of WT and miR-146a deficient mice. Histological assessment of bone microarchitecture paralleled our data obtained in μCT analysis (Sup. Fig. 3 A-D). Although overall numbers of osteoclasts (N.OC) were elevated in 6 and 12 months old miR-146a^−/−^ animals compared to WT animals, we could not detect a difference in osteoclast number per bone perimeter (N.OC/BPm) or osteoclast surface per bone surface (OC.S/BS) at all earlier time points up to 6 months. However, 12 months old mice deficient in miR-146a displayed reduced OC.S/BS and N.OC/BPm (Fig. 2 C-E).

**Fig. 3.**
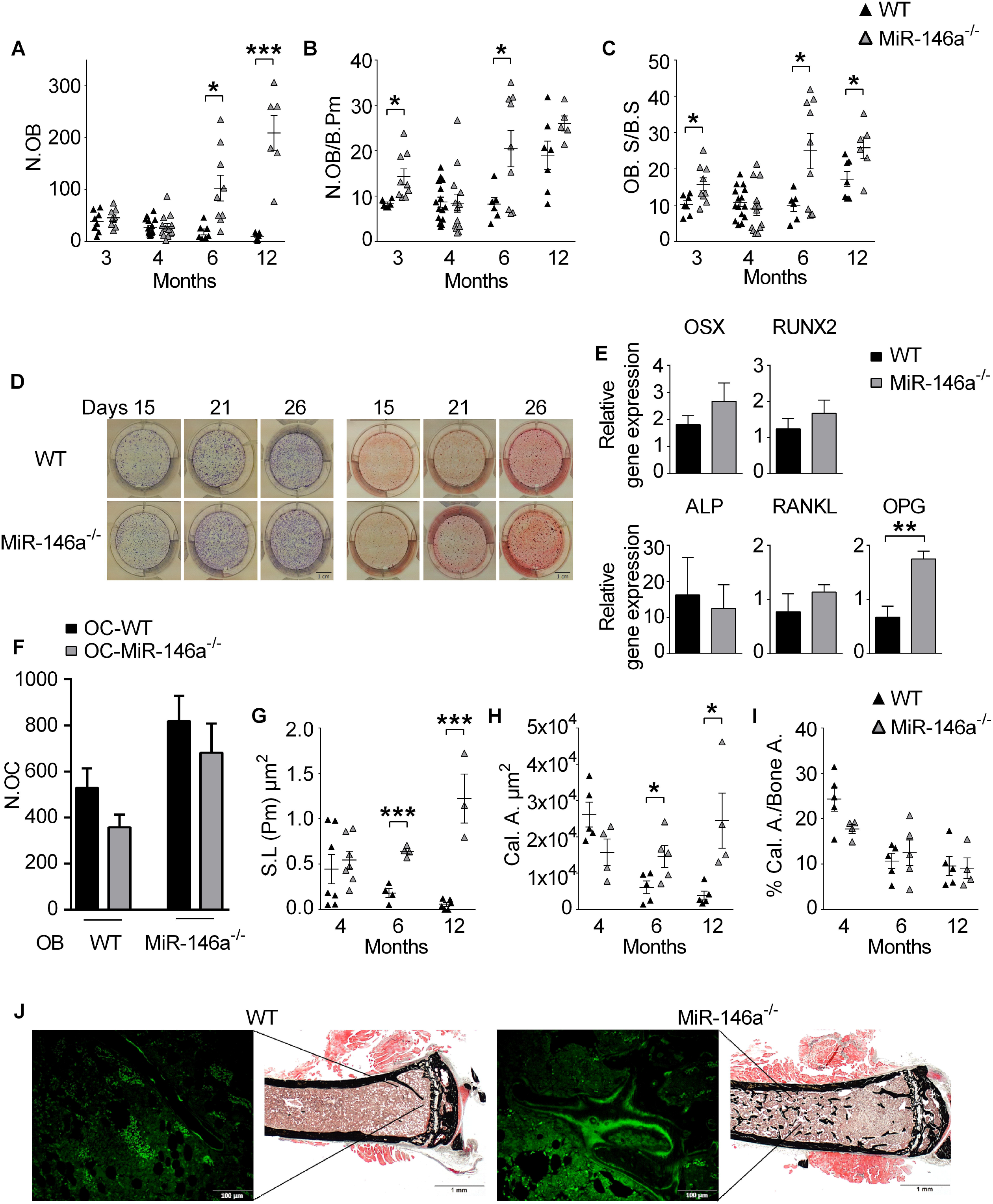
Increased activity of osteoblasts *in vivo* in aged miR-146a deficient mice. **A-C** Histomorphometric analysis of N.OB, N.OB/B.Pm and OB. S/BS of WT and MiR-146a^−/−^ mice aged 3 to 12 months of TRAP stained tibial sections (n≥5). **D** Representative images of alkaline phosphatase (left) and alizarin red (right) staining of WT and MiR-146a^−/−^ osteoblasts after 15, 21 and 26 days of osteogenic differentiation (bars 1cm, n=4). **E** Gene expression of OSX, RUNX2, ALP, RANKL and OPG was measured in osteoblasts of WT and MiR-146a^−/−^ mice after 21 days of osteogenic differentiation using Q-PCR (n≥3). **F** Numbers of osteoclasts were analyzed in co-cultures of either WT or MiR-146a^−/−^ osteoblasts cultured with WT or MiR-146a deficient bone marrow cells, stimulated with vitamin D3 and dexamethasone for 7 days (n=3). **G-I** WT and MiR-146a^−/−^ animals aged 4 to 12 months were labelled with calcein at two time points (6 and 1 day before sacrifice), histological sections of the proximal tibia were analyzed for S.L., Cal.A and Cal. A/Bone A. (n≥4). **J** Representative imagesv of on Kossa (left, bars 1mm magnification x 2,5) and calcein labelled (right, bars 100μm magnification x 20) tibial sections of WT and MiR-146a^−/−^ animals 6 months of age. N.OB, number of osteoblasts; N.OB/B.Pm, number of osteoblasts per bone perimeter; OB.S/BS, osteoblast surface per bone surface; OSX, osterix; ALP; alkaline phosphatase; RANKL, Receptor Activator of NF-κB; OPG, osteoprotegerin; S.L, single label; Cal.A, calcein area; Cal.A/Bone A, calcein area per bone area; All data shown were generated from female animals *p < 0.05; **p < 0.01; ***p < 0,001 Results are shown as mean ± SEM.

To explore if osteoclast differentiation and function are changed in mice lacking miR-146a we performed analysis of WT and miR-146a^−/−^ osteoclasts. Already published investigation of *in vitro* differentiated osteoclasts upon stimulation with RANKL had revealed no difference in bone marrow derived osteoclast numbers between WT and miR-146a deficient animals [8]. We extended these analyses to examine expression levels of miR-146a during osteoclastogenesis, which significantly increased initially after stimulation with RANKL and dropped back to baseline levels in mature OCs (Fig. 2 F). The tumor necrosis factor receptor-associated factor 6 (TRAF6), had been shown to be essential in RANKL-induced osteoclast generation, as TRAF6 is pivotal in RANK-mediated signal transduction and also to be a direct target of miR-146a [5, 19]. However, gene expression analysis of RANK and TRAF6 during *in vitro* differentiating osteoclasts did not show any difference between WT and miR-146a knock out animals, suggesting no regulation of TRAF6 levels by miR-146a during *in vitro* differentiating OCs (Fig. 2 F). Moreover, the function of *in vitro* differentiated OCs was not altered, as bone resorption capacity was not different between the two groups (Fig. 2 G). In addition, TRAF6 expression levels in bone measured by qPCR were not different at all timepoints analyzed, as were mRNA levels of SMAD4, another prominent experimentally confirmed target of miR-146a (Sup. Fig. 2 C).

To investigate whether deregulation of TRAF6 *in vivo* was responsible for the bone phenotype we observed in miR-146a deficient mice, we analyzed WT, miR-146a^−/−^ and miR-146a^−/−^/TRAF6^+/−^ animals. These miR-146a^−/−^/TRAF6^+/−^ mice were shown to express TRAF6 levels similar to WT animals, leading to the resolution of some important phenotypic observations in miR-146a deficient mice, such as aberrant myeloproliferation, splenomegaly and excessive inflammatory responses, but not others [6]. However, there were no significant differences in the BV/TV or other parameters analyzed in μCT between miR-146a^−/−^ and miR-146a^−/−^/TRAF6^+/−^ mice, neither in trabecular nor in cortical bone (Fig. 2 H-M and Sup. Fig. 4 A-C), demonstrating that deregulation of TRAF6 in miR-146a deficient mice is not responsible for the observed bone phenotype. Taken together, these data suggest that bone accrual in miR-146a deficient mice starting at 6 months of age was not due to decreased activity or numbers of bone resorbing osteoclasts.

**Fig. 4.**
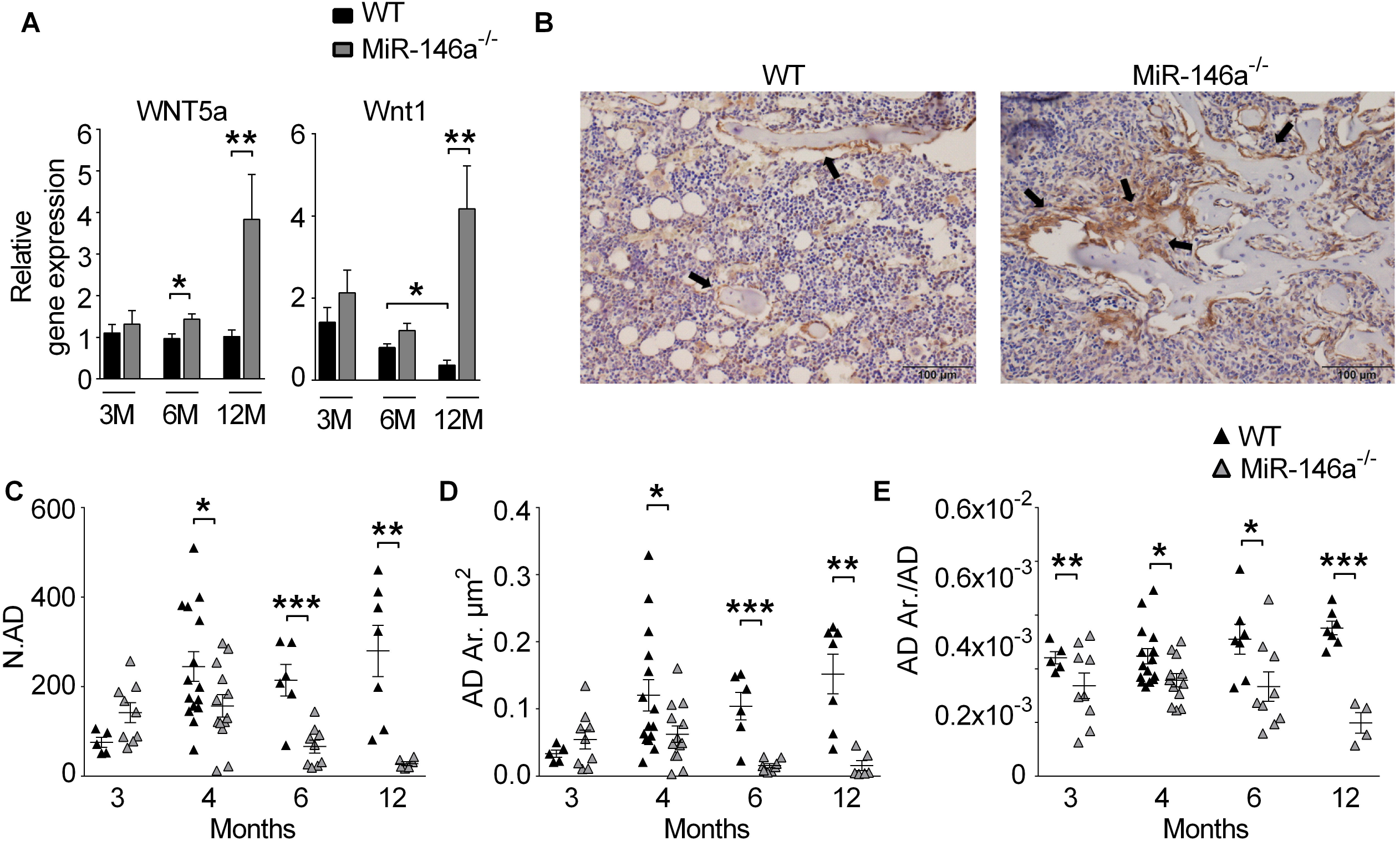
Loss of miR-146a increases Wnt signaling and prevents bone marrow adiposity with age. **A** Gene expression analysis of Wnt5a and Wnt1 in femoral bones of 3 to 12 months old WT and MiR-146a^−/−^ mice using Q-PCR (n≥6). **B** Representative pictures of immunohistochemically stained tibial sections of 6 months old WT and MiR-146a deficient animals using β-Catenin antibody (bars 100μm, magnification x 20). Black arrows indicate positively stained osteoblasts at sites of bone formation. **C-E** Histomorphometric analysis of N.AD, AD. Ar, AD. Ar/AD from histological sections of 3 to 12 months aged WT and MiR-146a^−/−^ animals (n≥4). N.AD, number of adipocytes; AD.Ar, adipocyte area; AD.Ar/AD, adipocyte area per adipocyte; All analysis shown were obtained from female animals *p < 0.05; **p < 0.01; ***p < 0,001 Results are shown as mean ± SEM.

### Increased activity of osteoblasts in vivo in aged miR-146a deficient mice

Analyzing osteoblast related parameters, we detected significantly increased numbers of osteoblasts per bone perimeter (N.OB/B.Pm) and osteoblast surface per bone surface (OB.S/BS) in miR-146a deficient mice, at 3, 6 and 12 months of age compared to their WT counterparts (Fig. 3 A-C). However, investigation of *in vitro* differentiation of calvarial osteoblasts did not reveal any difference in ALP staining, nor changes in the ability of OBs to induce bone nodule formation between WT and miR-146a^−/−^ animals, suggesting no major intrinsic defects in osteoblast differentiation *in vitro* (Fig. 3 D). In line with this observation, osteoblast related marker genes such as Runx2, Osterix or ALP did not show any significant differences between *in vitro* differentiated OBs of the two groups, only OPG was significantly increased in miR-146a deficient OBs compared to WT OBs (Fig. 3 E). To investigate whether OBs lacking miR-146a exhibit differences in their capacity to induce osteoclastogenic differentiation of bone marrow cells we performed co-culture assays. Calvaria-derived mesenchymal stem cells were cultured with bone marrow cells as myeloid responders, stimulated with vitamin D3 and dexamethasone. WT osteoblasts induced robust generation of osteoclasts, regardless of the genotype of the myeloid responder cells. Osteoblasts lacking miR-146a were similarly capable of stimulating osteoclastogenesis, again regardless of the genotype of the responder cells (Fig. 3 F).

To examine dynamic bone formation, we performed calcein labelling in WT and miR-146a deficient mice. While in WT animals single- or double-layer labelled bone surfaces were easily discernible, irregular incorporation of calcein occurred in miR-146a deficient animals, which precluded standard analysis of bone formation and mineral apposition rate. Therefore, we quantified total calcein incorporation in WT and miR-146a deficient mice. We detected significantly increased single layer calcein incorporation (Fig. 3 G) as well as an elevated total area of calcein incorporation (Fig. 3 H-J) in miR-146a deficient mice compared to WT mice starting at 6 months of age, indicating enhanced total activity of osteoblasts in miR-146a deficient mice.

### Increased Wnt expression and signaling in aged miR-146a deficient mice

Previous reports have demonstrated that miR-146a directly targets members of the Wnt family of proteins such as Wnt1, Wnt3 and Wnt5a [20, 21]. Wnt5a and Wnt1 were of particular interest, as mice deficient in these proteins have low bone mass due to reduced activity of osteoblast [22–24]. Analyzing RNA expression in bone, we detected similar levels of Wnt5a in bone tissue of mice aged 3 months. However, Wnt5a levels in bone were significantly increased in mice lacking miR-146a aged 6 months or older. In addition, while there was a trend towards increased expression of Wnt1 at 3 and 6 months of age, 12 months old mice displayed significantly increased expression of Wnt1 (Fig. 4 A). These data demonstrate, that Wnt family members, especially Wnt5a, are increased in aged but not in young miR-146a deficient mice. Wnt signaling, induced both by Wnt5a and Wnt1, leads to increased stability of the transcription factor β-Catenin and therefore increased presence of this protein [25, 26]. In line with augmented Wnt signaling, we detected markedly elevated levels of β-Catenin in bone tissue, and specifically in osteoblasts of aged miR-146a deficient mice compared to WT mice (Fig. 4 B arrows), demonstrating increased Wnt signaling in bone of miR-146a deficient mice compared to WT mice.

Wnt5a in particular was shown to be involved regulating differentiation of mesenchymal stem cells, where it drives osteogenesis at the expense of adipogenesis [27]. Since we had observed increased numbers of osteoblasts in aged miR-146a deficient mice, we next investigated bone marrow adiposity in those mice. Indeed, we found significantly increased numbers of adipocytes in WT mice compared to miR-146a deficient mice, suggesting increased osteoblast generation accompanied by decreased adipocyte generation and bone marrow adiposity during aging in miR-146a deficient mice compared to WT mice (Fig. 4 C-E).

### MiR-146a deficiency protects from ovariectomy induced bone loss

To analyze the mechanical properties of bone from miR-146a deficient animals, we performed 3-point bending assays. Bone strength of mice aged 6 months was not altered compared to control animals, and we detected an equal amount of collagen in bone of 6 months old WT and miR-146a^−/−^ animals (Fig. 5 A and B).

**Fig. 5.**
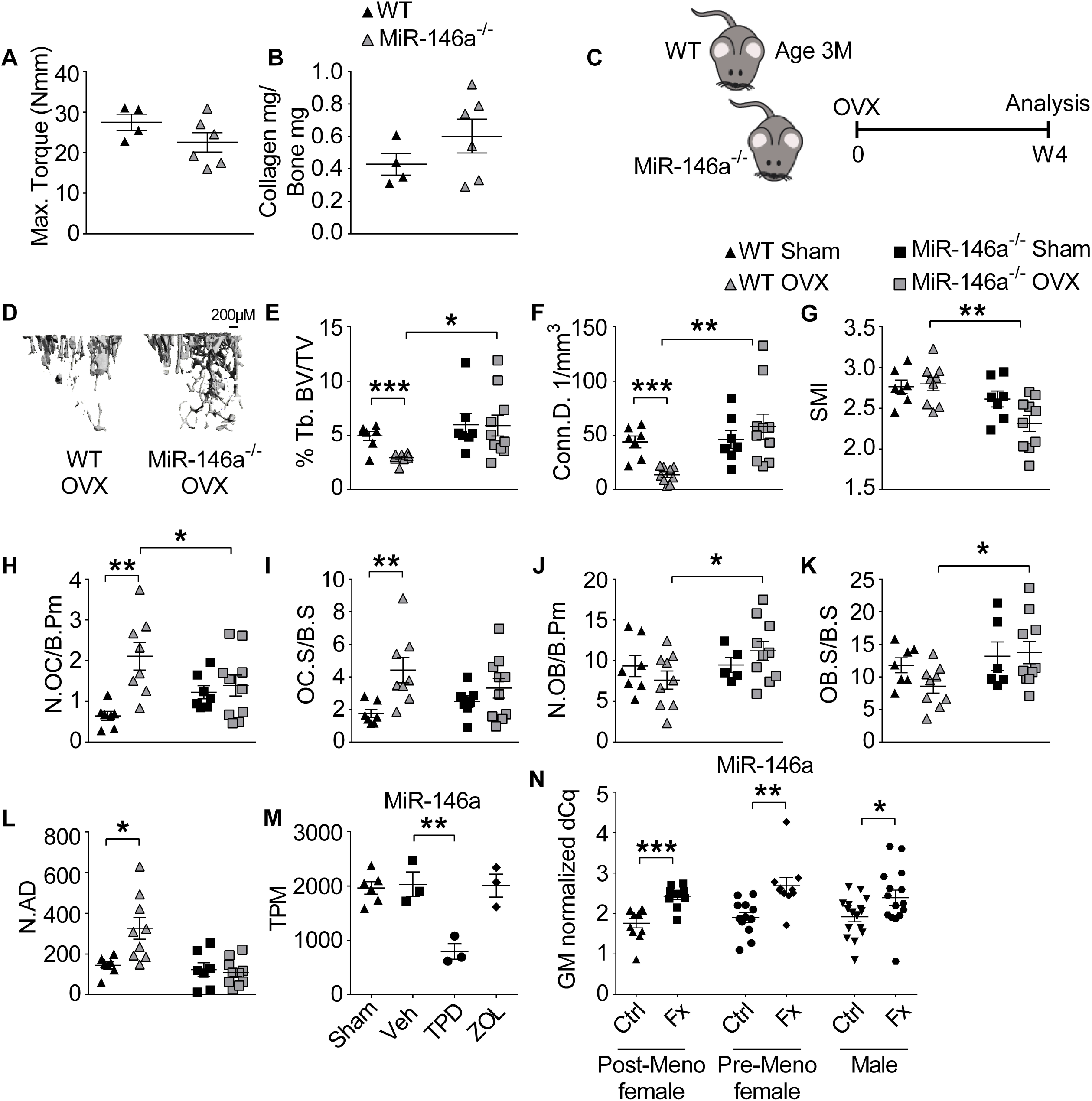
Loss of MiR-146a protects from ovariectomy-induced bone loss and is increased in patients with fragility fractures. **A** and **B** Maximal torque at the femoral shaft assessed by three point bending and Collagen mg/Bone mg analyzed in femoral bone in 6 months old WT and MiR-146a^−/−^ animals (n≥4). **C** Schematic illustration of the ovariectomy induced bone loss experiment performed in 3 months old WT and MiR-146a^−/−^ animals. **D** Representative μCT images of tibial trabecular bones from ovariectomized WT and MiR-146a deficient mice (bar 200μm). **E-G** Three-dimensional reconstruction and quantitation of Tb. BV/TV, Conn.D. and SMI of the trabecular tibial bone from sham as well as ovariectomized WT and MiR-146a^−/−^ mice using μCT (n≥7). **H-L** Histomorphometric analysis of N.OC/B.Pm, OC.S/BS, N.OB/B.Pm, OB.S/BS and N.OB from TRAP stained sections of tibial bone (n≥5). **M** MiR-146a levels were analyzed in sera of 6 months old sham operated or ovariectomized rats treated with either vehicle, TPD or ZOL starting 8 weeks after ovariectomy for 12 weeks, using Q-PCR (n≥3). **N** Serum levels of MiR-146a were measured in female post-menopausal, pre-menopausal and in male patients, control or low traumatic, using Q-PCR (n≥10). Tb. BV/TV, trabecular bone volume per tissue volume; Conn.D., connectivity density; SMI, structural model index; N.OC/B.Pm, numbers of osteoclasts per bone perimeter; OC.S/BS, osteoclast surface per bone surface; N.OB/B.Pm, numbers of osteoblasts per bone perimeter; OB.S/BS, osteoblast surface per bone surface; TPD, teriparatide; ZOL, zoledronate; TPM, total count per million; FX, fracture; GM, global mean (mean Cq-value); All analysis shown accept O, were obtained from female animals *p < 0.05; **p < 0.01; ***p < 0,001 Results are shown as mean ± SEM.

To evaluate the role of this miRNA in osteoporosis development, we used the murine ovariectomy (OVX)-induced model of postmenopausal osteoporosis. We performed OVX in 3 months (12 weeks) old WT and knock out animals, sham operated animals were used as controls. In line with our previous analyses, baseline μCT data showed no differences in bone parameters at this stage of aging (Fig. 5D). After 4 weeks, ovariectomized WT animals showed robust bone loss compared to sham operated mice. In contrast, OVX in miR-146a deficient mice did not result in any detectable bone loss compared to sham operated miR-146a deficient mice (Fig. 5 C-G and Sup. Fig. 5 A-G). Moreover, histological analysis of bone resorbing and forming cells showed elevated numbers and activity of osteoclasts, as determined by increased number of osteoclasts per bone perimeter and osteoclast surface per bone surface, in WT OVX animals compared to sham operated mice. In contrast, miR-146a deficient animals showed no change in osteoclast numbers and activity (Fig. 5 H and I). On top of that, osteoblast numbers per bone perimeter and osteoblast surface per bone surface even increased in OVX knock out animals compared to WT animals (Fig.5 J-K), suggesting increased activity of osteoblasts in miR-146a deficient compared to WT mice after OVX. In addition, while bone marrow adiposity measured by the number of adipocytes increased significantly in WT mice after OVX, we detected no change in miR-146a deficient mice after OVX (Fig. 5L). Summarized, these data reveal reduced activation of osteoclast and preserved activity of osteoblasts in ovariectomized miR-146a deficient mice, leading to complete protection from OVX-induced bone loss.

In order to investigate how current treatment regimens for estrogen deficient bone loss influence miR-146a expression in bone tissue, we reanalyzed data from an OVX rat model in wild type animals, using sham operated animals as controls. After eight weeks, OVX animals received either vehicle (Veh), anti-resorptive treatment using zoledronate (ZOL) or osteo-anabolic treatment using teriparatide (TPD), a recombinant parathyroid hormone (PTH) shown to increase bone mass of rats, for 12 weeks [28]. Analyzing microRNA expression in bone, we found that, in line with our hypothesis, treatment with TPD significantly reduced the levels of miR-146a in bone tissue, correlating with an almost complete protection from OVX induced bone loss. In contrast, treatment with bisphosphonates which target osteoclast-mediated bone resorption, did not alter levels of miR-146a in bone. (Fig. 5 M). These data demonstrate that treatment of OVX-induced bone loss with recombinant PTH is associated with suppression of miR-146a.

### Increased expression of miR-146a in patients with fragility fractures

Finally, in order to investigate whether miR-146a could also be involved in human disease, we tested the levels of miR-146a in patients with osteoporosis, reanalyzing an existing dataset [29]. In line with our murine data, women suffering low trauma fractures had increased circulating serum levels of miR-146a compared to those who did not (Fig. 5 N). There was even a significant difference in pre-menopausal fractures in females compared to controls. In men, miR-146a was significantly elevated in individuals who suffered a fracture as well. However, the difference was less pronounced in males compared to females, pointing towards a predominant role of miR-146a in the regulation of bone in female individuals.

## Discussion

In this study, we identify a novel and unexpected role of miR-146a in bone homeostasis. MiR-146a increases with age and acts as an epigenetic molecular switch terminating bone accrual, leading to bone loss during aging by downregulating bone anabolic Wnt signaling. Conversely, loss of miR-146a leads to continuous bone accrual during aging as a result of unrestricted bone anabolism and intact Wnt signaling. Therefore, miR-146a crucially controls the temporal facet of bone homeostasis.

MiR-146a has been primarily identified as an anti-inflammatory miRNA, as it has been shown to inhibit many aspects of inflammation, hematopoetic stem cell biology and cancer [6, 7, 9, 30–32]. However, miR-146a has also been implicated in immune dysfunction during aging, as it has been shown to accumulate in macrophages and dendritic cells of aged mice, leading to dysfunction of these cells [10, 33].

Moreover, miR-146a was found to suppress the osteogenesis of adipose derived mesenchymal stem cells by targeting SMAD4 [34]. Since SMAD4 was not differentially expressed in our analysis of miR-146a deficient bone, neither in young nor aged animals it does not seem to be responsible for the elevated bone growth in knock out animals.

The role of Wnt proteins and their receptors in bone biology is well established [25, 26]. The role of Wnt5a is of particular interest, as it has been implicated in osteoclast activity as well as osteoblast differentiation and activity. The net effect in full Wnt5a knock out mice is reduced bone mass, accompanied by severely reduced activity of osteoblasts [22]. Loss of Wnt5a specifically in osteoclasts leads to reduced bone mass due to decreased bone formation, whereas deletion of its receptor Ror2 in OCs leads to increased bone mass, suggesting that Wnt5a activates both osteoblasts and osteoclasts, with a predominant effect on osteoblast function [22, 35]. Mutations in Wnt1 have been shown to be responsible for osteoporosis and osteogenesis imperfecta, and transgenic overexpression of Wnt1 has been shown to markedly increase bone mass in mice [4, 36]. In this study we found deregulation of Wnt5a as well as Wnt1 accompanying the increased osteoblast activity in miR-146a deficient mice. Importantly, young mice show similar levels of Wnt5a and Wnt1 in WT and miR-146a deficient mice, demonstrating that Wnt5a is temporally regulated, as it is the case with miR-146a. The fact that timing of Wnt5a expression is crucial for some of its effects has also been demonstrated in embryogenesis, with the help of tamoxifen inducible Wnt5a expression in mice [37]. Wnt proteins have been shown to decline with age in bone [38, 39]. We also find a decline in Wnt1 in our analyses in WT mice, but not in miR-146a deficient mice. In addition, classical, but also sometimes non-classical Wnt signaling leads to stabilization of β-Catenin [40]. Increased stabilization of β-Catenin in osteoblasts has been shown to increase bone mass, whereas deletion leads to reduced bone mass [41]. In our experiments, aged miR-146a deficient mice exhibit markedly increased presence of β-Catenin in bone, as a result of increased Wnt signaling. The effects of miR-146a deficiency are unique, as this miRNA is controlling the temporal aspect of bone turnover. This contrasts with mice deficient in sclerostin, an endogenous inhibitor of Wnt signaling, leading to a high bone mass phenotype already at a younger age [42]. Interestingly, miR-146 is increased in senescent cells [43], whose elimination prevents bone loss with aging [44].

Wnt signaling has also been implicated in regulating the fate of mesenchymal stem cells to differentiate into either the osteoblastic or the adipogenic lineage, favoring the former over the latter [27, 45]. In WT mice, bone marrow adiposity continuously increases with age. In line with increased activity of Wnt proteins, we detected even decreasing bone marrow adiposity and increasing numbers of osteoblasts over time in miR-146a deficient mice. In addition, miR-146a deficient mice are, in accordance with a recent report [46], protected from OVX-induced bone loss. Interestingly, also during ovariectomy, we detected increased osteoblast and reduced generation of adipocytes in miR-146a deficient mice. As bone marrow adiposity has been shown to increase with age in humans and has been linked with osteoporosis in the elderly as well [47], we propose that miR-146a favors adipocyte generation at the expense of osteoblasts during aging, leading to reduced osteoanabolic capacities as a result of reduced osteoblast differentiation.

Therefore, the temporal regulation of miR-146a expression in bone seems to provide the molecular clock, which restricts osteoblast generation by decreasing Wnt5a and Wnt1 expression, tipping the scale towards bone loss during aging in WT mice. This important regulatory function is lost in miR-146a deficient mice, leading to continuous bone accrual during aging. Importantly, levels of miR-146a are increased in the circulation of patients suffering from post-menopausal fragility fractures. Therefore miR-146a potentially plays an important role in human osteoporosis.

Therefore, our study describes a novel molecular checkpoint controlling age related bone loss and could lead to therapeutic targeting of miR-146a for osteoporosis and age-related bone loss.

## Materials und Methods

### Mice

Breeding pairs of miR-146a^−/−^ and miR-146a^+/+^ littermate (B6.(FVB)-MIR146 TM1.1BAL/J) mice were provided by Mark Boldin/David Baltimore. Bones from miR-146a^−/−^ and miR-146a^−/−^/TRAF6^+/−^ mice were obtained from M.Boldin. Mice used in all experiments were age- and sex-matched. All data were generated from littermates.

### Analysis of bone parameters

Histomorphometry and bone formation analysis of mouse tibia from WT, miR-146a^−/−^ was analyzed as described [8]. For von Kossa stainings tibias were fixed in paraformaldehyde at room temperature for 6 hours. Bones were dehydrated using a gradual series of ethanol (70%, 95%, and 100%), infiltrated and embedded without decalcification in methylmethacrylate and von Kossa staining was done on tibial sections. For μCT analysis we used SCANCO Medical μCT 35 to produce images from trabecular and cortical bone and analyzed with SCANCO evaluation software for segmentation, three-dimensional morphometric analysis, density and distance parameters.

### Immunohistochemisty

For immunohistochemistry β-Catenin (BD Bioscience) at a dilution of 1:200 was used.

### Cell culture

Osteoclast generation was done as previously described [8].

For bone resorption assays osteoclasts were seeded 600 000 cells/well on osteo assay surface plates (Corning) and cultured as previously described [8]. After 4 days osteoclasts were removed using 5% Sodiumhypochlorid for 5 minutes and washed to times with water, water was removed, and wells were air dried.

Osteoblasts were isolated from calvariae of neonatal WT and miR-146a^−/−^mice. Calvariae were digested for 10 minutes in alpha MEM (Gibco) containing 0,1% Collagenase (Sigma) and 0,2% DispaseII (Sigma). Osteogenic precursors were expanded in alpha MEM containing 10% FCS (Gibco) and 1% penicillin /streptomycin. Differentiation of osteoblasts were achieved by adding 0,2mM L-Ascorbate and 10mM β-Glycerophosphate (both Sigma). After 15, 21 and 26 days, cells were fixed with 4% formalin, following staining with either aqua bidest containing 8% alizarin red (Sigma) or staining for alkaline phosphatase using 5-Brom-4-chlor-3-indoxylphosphat/ Nitro blue.

### MiRNA measurement

Micro RNA was isolated using miRNeasy Mini Kit (Qiagen). MiR-146a expression was measured using TaqMan miRNA Assays hsa-miR-146a (Applied Biosystems) according to the manufacturer’s instruction using the Rotor-Gene Q PCR cycler (Qiagen). U6snRNA (miRNA assay U6 snRNA Applied Biosystems) was used as internal control. Relative expression of miR-146a was calculated by the 2ΔΔCT method.

### Quantitative real time PCR

For mRNA expression analysis, total RNA was isolated from OCs, OBs and minced femur using RNeasy mini kit (Qiagen). CDNA was prepared using the omniscript RT kit (Qiagen), followed by SBR green-based quantitative PCR (Roche) using the Light cycler 480 (Roche). MRNA amounts were normalized relative to glyceraldehyde-3-phosphatedehydrogenase (GAPDH) mRNA. Generation of the correct size amplification products was confirmed with agarose gel electrophoresis. The primers for real time PCR were as follows: *RANK*: 5’-CACAGACAAATGCAAACCTTG-3’ and 5’-GTCTTCTGGAACCATCTTCCTCC-3’; *TRAF6: 5’-* AAAGCGAGAGATTCTTTCCCTG-3’ and 5’-ACTGGGGACAATTCACTAGAGC-3’; *OSX:* 5’-GGAGGCACAAAGAAGCCATAC-3’ and 5’-TGCAGGAGAGAGGAGTCCATTG-3’; *RUNX2:* 5’*-* TGGCTTGGGTTTCAGGTTAGGG-3’ and 5’-TCGGTTTCTTAGGGTCTTGGAGTG-3’; *ALP:* 5’*-* GCTGATCATTCCCACGTTTT-3’ and 5’-CTGGGCCTGGTAGTTGTTGT-3’; *RANKL*: 5’-TCGTGGAACATTAGCATGGA-3’ and 5’-CCTCTCCCAATCTGGTTCAA-3’; *OPG*: 5’-TACCTGGAGATCGAATTCTGCTT-3’ and 5’-CCATCTGGACATTTTTTGCAAA-3’; *Wnt5a*: 5’-CCAACTGGCAGGACTTTCTC-3’ and 5’-GCATTCCTTGATGCCTGTCT-3’; *Wnt1*: 5’-TTTTGGTCGCCTCTTTGG-3’ and 5’-TGCCTCGTTGTTGTGAAGG-3’; *OC*: 5’-ACCTTATTGCCCTCCTGCTT-3’ and 5’-GCGCTCTGTCTCTCTGACCT-3’; *OPN*: 5’-CTCCATCGTCATCATCATCG-3’ and 5’-TGCACCCAGATCCTATAGCC-3’; *TRAP*: 5’-TCCTGGCTCAAAAAGCAGTT-3’ and 5’-ACATAGCCCACACCGTTCTC-3’; *CatK*: 5’-TGAGAGTTGTGGACTCTGTGCT-3’ and 5’-TTGTGCATCTCAGTGGAAGACT-3’; *SMAD4*: 5’-TGGGTCCGTGGGTGGAATAG-3’ and 5’-TCTAAAGGCTGTGGGTCCGC-3’; GAPDH: 5’-TGGCATTGTGGAAGGGCTCATGAC-3’ and 5’-ATGCCAGTGAGCTTGCCGTTCAGC-3’. The relative expression of the mRNA of the gene of interest was calculated by the 2ΔΔCT method.

### Dynamic labelling of bone

Calcein labelling was performed and analyzed as described [48].

### Ovariectomy (OVX)

OVX of 3 months old female WT and miR-146a^−/−^ animals was performed as described [49]. Ovaries were removed from OVX animals, skin of sham operated female animals was incised, ovaries were left intact. We histological analysis of surgically removed ovaries was performed to verify OVX. After 4 weeks of recovery and to allow the onset of osteoporosis, all mice were sacrificed. Bone loss was evaluated in the tibiae, of these mice, histologically and using μCT image analysis.

### Three-point bending and mechanical stability

Mechanical properties of femur from WT and miR-146a^−/−^ (6 months old animals) were tested as described [50].

### ELISA

Serum concentration of RANKL (R&D Systems), CTXI (Biozol/Biomedica), OPG (AbFrontier), P1NP (ids immunodiagnosticsystems), Estradiol (Calbiotech) were assessed according to the manufacturer’s instructions.

### Collagen amount

For collagen determination femurs of 6 months old WT and miR-146a−/− mice were taken. Bone marrow removed, and tissue was dissolved at a concentration of 100 mg/ml in 12 M HCl. Afterwards, the concentration of the hydroxyprolines, and consequently, of the collagen was determined using a Total Collagen Assay Kit (QuickZyme, Leiden Netherlands) with rat tail collagen type I as a standard, according to the manufacturer’s instructions.

### Statistical analysis

Statistical significance of two different groups was calculated using the unpaired two tailed Student t-test. All analyses were performed using GraphPad Prism 6 software. Graphs present data as mean ± SEM. A p value less than or equal to 0.05 was considered significant (*p < 0.05, **p < 0.01, ***p < 0.001).

## Acknowledgements

We thank Tetyana Shvets and Carl-Walter Steiner for expert technical assistance. This research has received funding from the Innovative Medicines Initiative 2 Joint Undertaking under Grant Agreement no 777357 (RTCure) and by the European Union Horizon 2020 research and innovation program under grant agreement 675228, the Lilly Innovation Award of the Austrian Society of Rheumatology and the ‘Cells-in-Motion Cluster of Excellence’ (EXC 1003–CiM), University of Münster, Germany. We also thank the Biomedical Sequence Facility of the Medical University Vienna for their support.

## Supplementary Figure Legends

**Sup Fig. 1**

**Male animals lacking miR-146a show a milder bone phenotype than female mice.**

Histomorphometric analysis of bone volume per tissue volume (BV/TV) TRAP stained tibial sections of male WT and MiR-146a^−/−^ animals aged 3 to 12 months (n≥5). Results are shown as mean ± SEM.

**Sup Fig. 2**

**MiR-146a regulates osteoblast and osteoclast gene expression during aging.**

Gene expression analysis of femoral bones from 3 to 12 months aged WT and MiR-146a^−/−^ animals. Osteoblast marker genes RUNX2, OC, ALP, OSX, OPN (**A**); Osteoclast marker genes TRAP, CTSK and RANK (**B**) and MiR-146a target genes SMAD4 and TRAF6 (**C**) were analyzed using Q-PCR (n≥4). Runx2, runt-related transcription factor 2; OC, osteocalcin; ALP, alkaline phosphatase; OSX, osterix; OPN, osteopontin; TRAP, tartrate resistant acid phosphatase; CTSK, cathepsin K; RANK, receptor activator of NF-κB, SMAD4, SMAD family member 4; TRAF6, tumor necrosis factor receptor-associated factor 6; All data shown were obtained from female animals. *p < 0.05; **p < 0.01; ***p < 0,001 Results are shown as mean ± SEM.

**Sup Fig. 3**

**Deficiency of MiR-146a leads to elevated trabecular bone growth.**

Histomorphometric analysis of TRAP stained tibial sections of WT and MiR-146a^−/−^ animals aged 3 to 12 months. BV/TV, Tb.Th, Tb.Sp and Tb.N are shown (n≥3). BV/TV, bone volume per tissue volume; Tb. Th, trabecular thickness; Tb. Sp, trabecular separation; Tb.N, trabecular number; Depicted data were generated from female animals *p < 0.05; **p < 0.01; ***p < 0,001 Results are shown as mean ± SEM.

**Sup Fig. 4**

**The MiR-146a target TRAF6 does not contribute to age dependent bone regulation mediated by this miRNA.**

**A-C** μCT analysis of trabecular bone from proximal tibias of 3 and 6-month-old WT, MiR-146a^−/−^ and MiR-146a^−/−^ TRAF6^+/−^ mice. Tb. N, trabecular number; Tb.Th, trabecular thickness and Tb.Sp, trabecular separation are depicted (n≥3). Data shown were generated from female animals. Results are shown as mean ± SEM.

**Sup Fig. 5**

**Loss of MiR-146a protects from ovariectomized induced bone loss.**

**A-G** μCT analysis of trabecular tibial bone from sham as well as ovariectomized WT and MiR-146a^−/−^ mice. Tb. N, trabecular number; Tb.Th, trabecular thickness; Tb.Sp, trabecular separation; Ct.BV/TV, cortical bone volume per tissue volume; Ct. Porosity, cortical porosity; Ct.Th, cortical thickness and BMD, bone mineral density are displayed (n≥5). Data were generated from female animals *p < 0.05; **p < 0.01; ***p < 0,001 Results are shown as mean ± SEM.

