## Supplementary figures and images for "MiR-146a controls age related bone loss"

### Supplemental Figures

**Sup. Fig. 1**

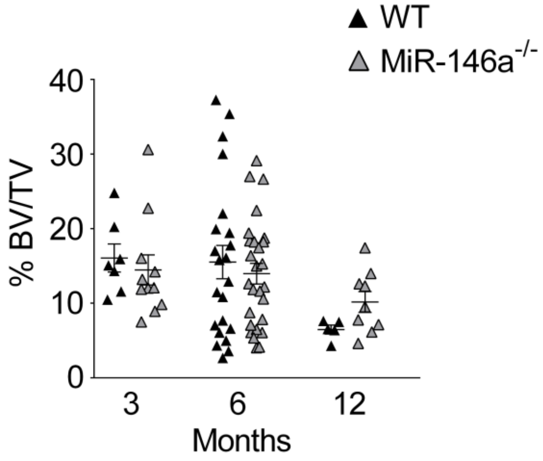

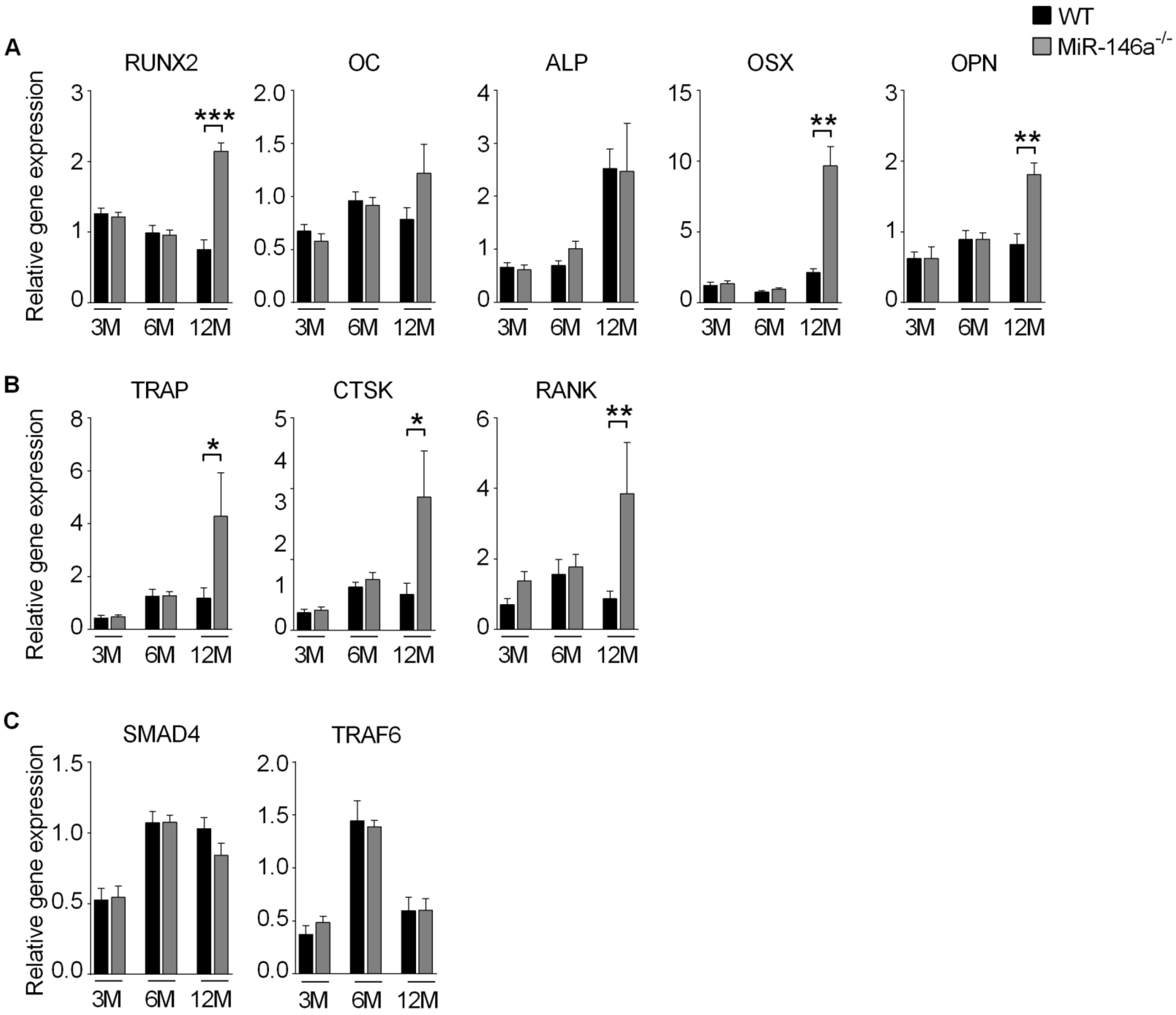

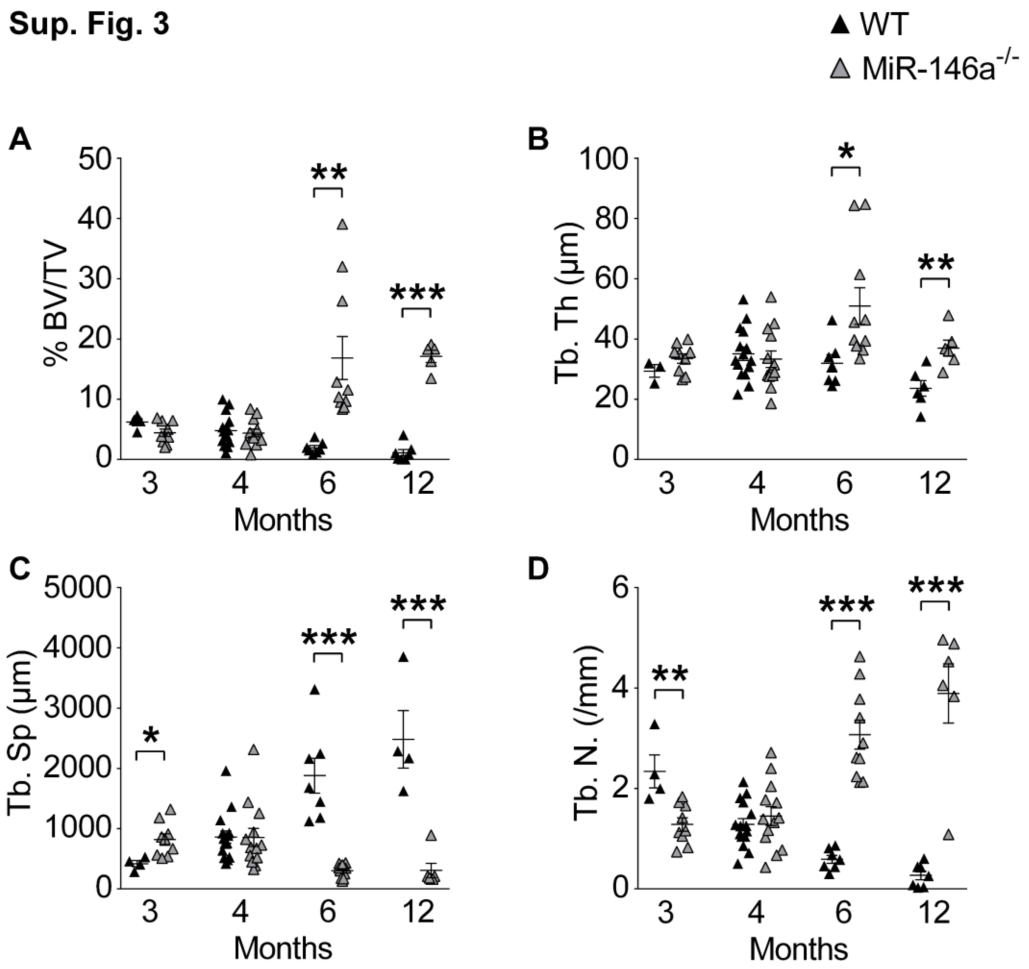

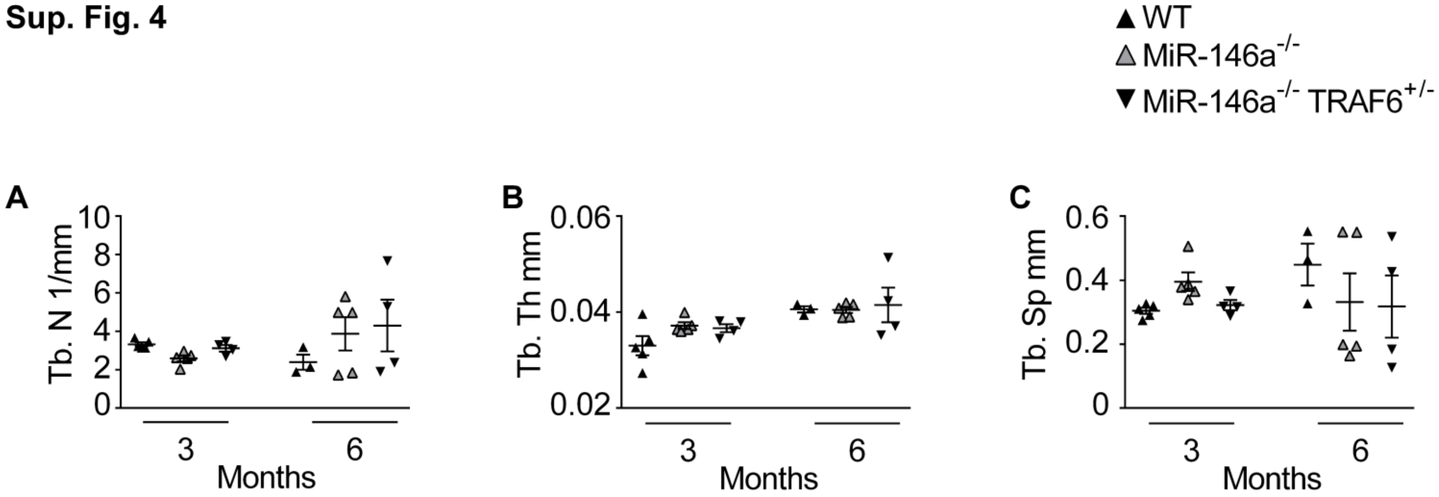

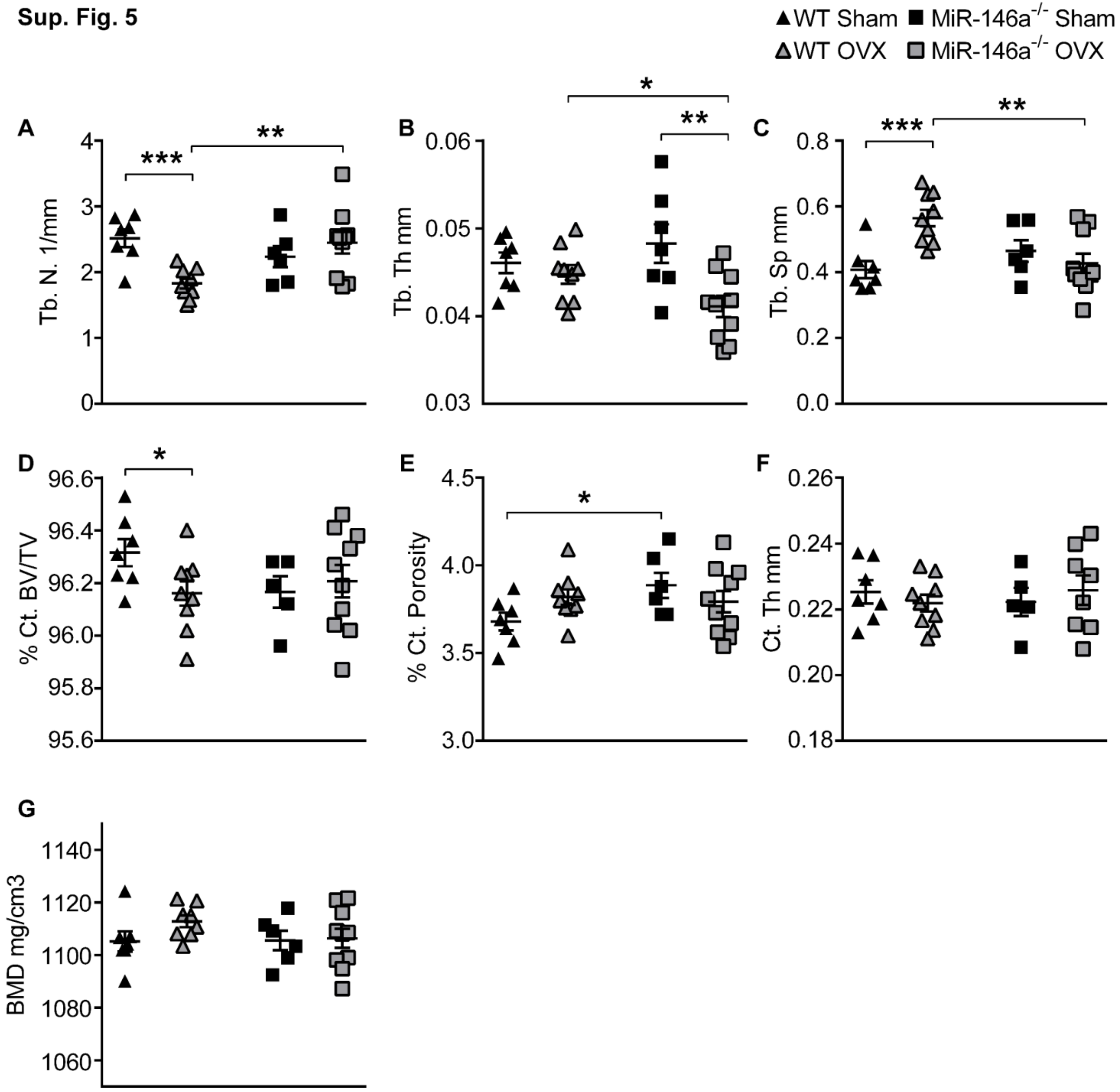
